# Leeches Predate on Fast-Escaping and Entangling Blackworms by Spiral Entombment

**DOI:** 10.1101/2024.05.14.594257

**Authors:** Harry Tuazon, Samuel David, Kenneth Ma, M. Saad Bhamla

## Abstract

We investigate how the *Helobdella* spp. freshwater leeches capture and consume *Lumbriculus variegatus* blackworms despite the blackworm’s ultrafast helical swimming escape reflex and ability to form large tangled ‘blobs’. We describe our discovery of a unique spiral ‘entombment’ strategy used by these leeches to overcome the blackworms’ active and collective defenses. Unlike their approach to less reactive and solitary prey like mollusks, where leeches simply attach and suck, *Helobdella* leeches employ this spiral entombment strategy specifically adapted for blackworms. Our findings highlight the complex interactions between predator and prey in freshwater ecosystems, providing insights into ecological adaptability and predator-prey dynamics.

## Predatory Behavior of Helobdella Leeches

Freshwater leeches, particularly *Helobdella* spp., are known for their diverse dietary habits, consuming a variety of prey ranging from oligochaetes to mollusks and insect larvae (Kutschera et al. 2013; Saglam et al. 2018, 2023). As apex predators in small ponds and lakes, they serve as models for exploring ecological and evolutionary themes, including interspecies competition and niche overlap (Govedich et al. 2010). *Helobdella* leeches, characterized by a pale gray or yellow hue, a flattened head with twin ocular spots, and a body length under two centimeters, display a fascinating biological profile (Fig. 1c, and inset). Unlike macrophagous leeches that ingest prey whole, liquidosomatophagous leeches like *Helobdella* draw nutrients from the bodily fluids of their prey using a specialized proboscis, showing a preference for oligochaetes, as evidenced by oligochaete proteins in their digestive systems (Kutschera et al. 2013; Sawyer 1972, 1986).

**Fig. 1.**
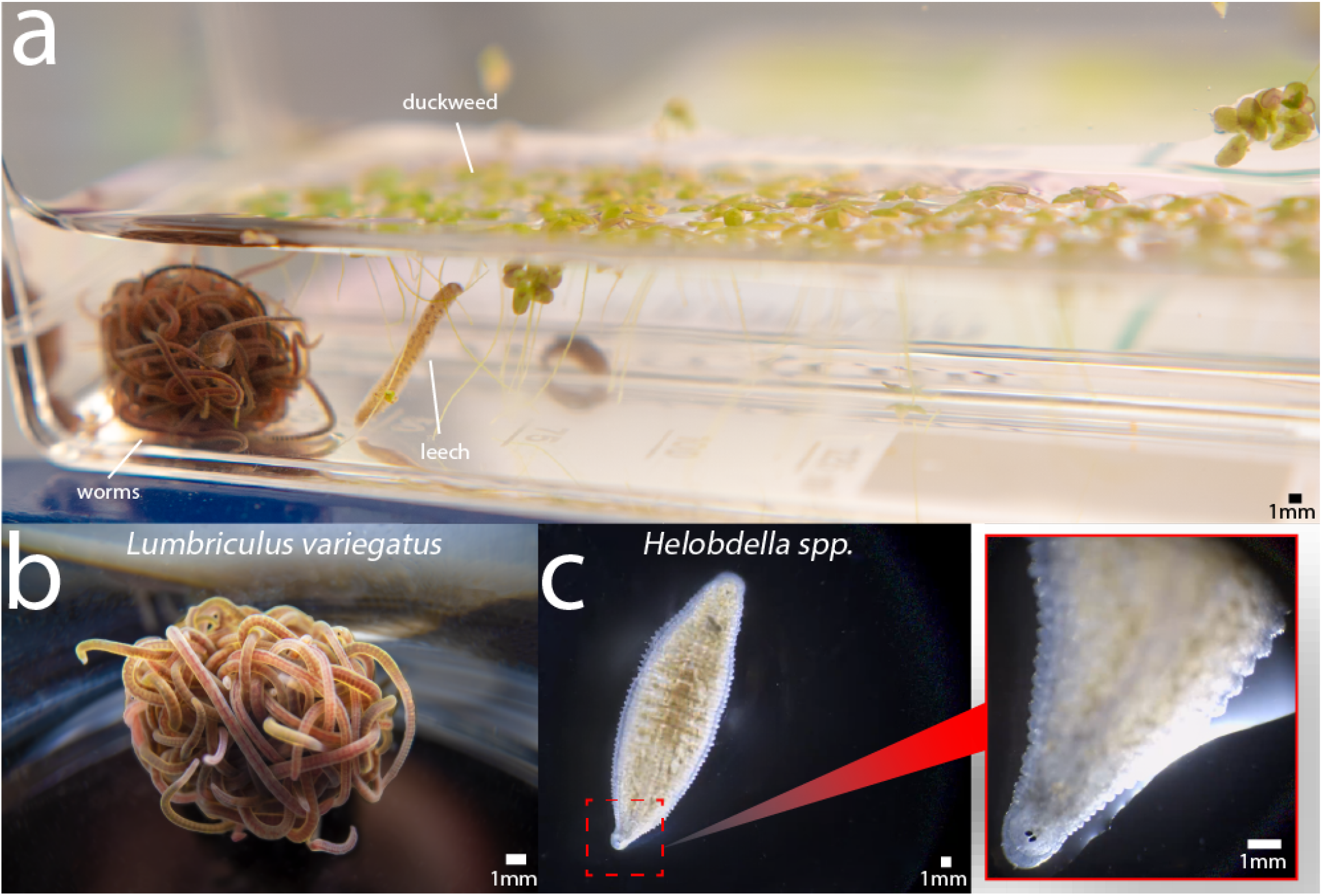
Interaction Between *Helobdella* spp. Leeches and *Lumbriculus variegatus* blackworms. **(a)** *Helobdella* spp. uses its anterior sucker to explore the area near the roots of duckweed and an entangled worm blob of N*∼*20 *Lumbriculus variegatus* blackworms. **(b)** The prey: California blackworms (*Lumbriculus variegatus*). **(c)** The predator: Freshwater leech: *Helobdella* spp. The inset shows the simple eyes of the leech.

The interaction between *Helobdella* leeches and California blackworms, *Lumbriculus variegatus*, exemplifies complex predator-prey relationships in freshwater ecosystems (Fig. 1a). *Helobdella* display advanced trapping and survival techniques beyond simple predation when targeting oligochaetes like the blackworm (Fig. 1b). Among oligochaetes, *Helobdella* show a preference for blackworms or *T. tubifex* (sludge worms), depending on habitat productivity levels (Young 1980; Young and Procter 1985).

Blackworms exhibit rapid escape mechanisms through a helical swimming gait and also form protective entangled collectives, or “worm blobs,” comprising up to thousands of conspecifics (Patil et al. 2023; Drewes and Fourtner 1989; Drewes 1990). This collective behavior, much larger in size than the leeches themselves, provides a unique perspective on natural selection pressures and biomechanical influences on predatory behaviors (Ozkan-Aydin et al. 2021; Patil et al. 2023; Tuazon et al. 2022, 2023). These robust escape reflexes give blackworms a survival advantage against predators like *Helobdella* leeches. Given these defenses, we ask: How do *Helobdella* leeches capture and consume blackworms despite the blackworms’ ultrafast helical swimming escape reflex and ability to form large entangled collectives?

In this study, we describe our discovery of a unique spiral ‘entombment’ strategy used by *Helobdella* leeches to overcome the blackworms’ active and collective defenses. Unlike their approach to less reactive and solitary prey like mollusks, where leeches simply attach and suck, *Helobdella* leeches employ this spiral entombment strategy specifically adapted for blackworms. We examine the behaviour strategy of this predatory action, exploring how these leeches manipulate their environment to encase and consume a worm. Our analysis reveals how the entombment process overcomes the dynamic defenses of worm blobs. Finally, we discuss the complex interactions between predator and prey in freshwater ecosystems, providing insights into ecological adaptability and predator-prey dynamics.

## Materials and methods

### Animals

We sourced California blackworms (*Lumbriculus variegatus*) from Ward’s Science, where they often came with freshwater leeches (*Helobdella spp*.) as unintended pests. Initially, we identified these leeches as *Helobdella stagnalis* based on general morphological features (Govedich et al. 2010). Later, more detailed morphological assessments as described by Saglam, et al. suggested they are more accurately as *Helobdella echoensis*, particularly due to the distinct positioning and shape of the eyes (Fig. 1c, inset) (Saglam et al. 2018). In the absence of comprehensive molecular analyses such as DNA or RNA sequencing for identification, we refer to these organisms broadly as *Helobdella* spp. or simply freshwater leeches throughout this paper.

We maintained the rearing conditions for the worms as described in our earlier publications (Tuazon et al. 2022, 2023). We isolated the freshwater leeches from the worms and stored them in refrigerated conditions. Before the experiments, we subjected the freshwater leeches to a starvation period of at least one week. Our research did not require approval from the Institutional Animal Care Committee, as studies involving invertebrate blackworms and freshwater leeches are exempt from these regulations.

### Data Acquisition

We recorded predator-prey interactions using a Leica MZ APO microscope (Heerbrugg, Switzerland) equipped with an ImageSource DFK 33UX264 camera (Charlotte, NC) at 30 FPS. The arena was a 35 mm petri dish containing filtered water. We randomly selected healthy worms (length 17.6±1.8 mm with both anterior and posterior segments) for the experiments. We chose freshwater leeches of similar size to the worms. Before introducing the prey, each leech spent one hour in the arena containing only water as a control period. We repeated each predation trial n=5 times.

## Spiral Entombment Predation Strategy of Helobdella Leeches

During predation of a single blackworm, *Helobdella* employs a unique hunting strategy, using its anterior sucker to latch onto a worm and swiftly envelop it in a spiral cocoon, illustrated in Fig. 2a. This “entombment” process often requires multiple attempts and involves meticulous coordination: finding an optimal area with the anterior sucker (t = 0 s), extending through flexure (t = 0 to 3 s), and executing coordinated ventral folding to form a cavity (t = 3 to 9 s). In this cavity, the leech inserts its the proboscis into the entombed worm to feed on its internal liquids and tissues through liquidosomatophagy (Fig. 2b) (Lynggaard et al. 2022).

**Fig. 2.**
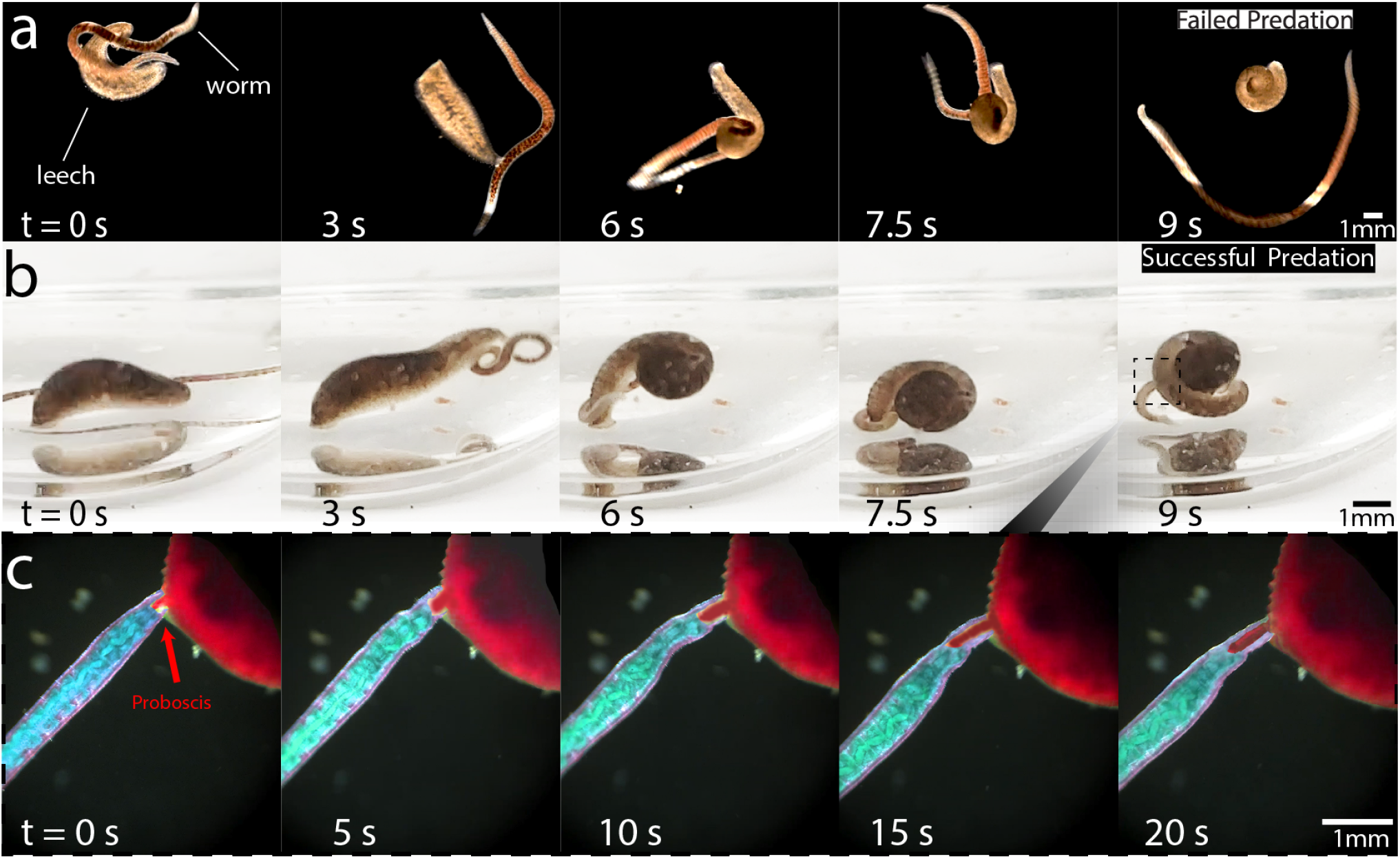
Predation strategy of *Helobdella* spp. against *Lumbriculus variegatus*. **(a)** Sequential frames of a failed predation attempt by a leech from the top perspectives. **(b)** A separate but successful predation attempt from the side view. The time stamps for both (a) and (b) are approximately similar. Initially, the leech secures a position along the worm’s body with its anterior sucker at t=0s and proceeds to stretch out at t=3s. In the ensuing entombment phase, spanning t=3s to t=7.5s, the leech performs a series of coordinated ventral folds coupled with a slight medial flexure, actions intended to form a cavity to entrap the worm. Failure to fully ensnare the prey often leads to its escape as shown in t=9s (a). If successfully trapped and immobilized as shown in t=9s (b), **(c)** the leech employs its proboscis (red arrow) to extract the internal contents of the worm. False coloring is added for visual clarity. (**Supplementary movie 1 and 2)**.

The spiral formation observed during this entombment provides biomechanical advantages such as enhanced grip and control, qualitatively similar to those seen in elephants, seahorse, chameleons, and various plants and flowers (Porter et al. 2015; Herrel et al. 2013; Takaki et al. 2003). This is complemented by a series of latching, releasing, and ventilating actions that facilitate respiration in the low-oxygen, detritus-rich environments typical of their habitats (Mann 1956; Milne and Calow 1990).

Both species, primarily found along the shallow, stagnant edges of freshwater systems in North America and Europe, exhibit adaptations to their environments that influence their interactions (Saglam et al. 2018; Kutschera and Weisblat 2015; Timm and Martin 2015; Govedich et al. 2010). *Helobdella* requires substrate attachment, via their posterior sucker, for successful feeding, indicating a reliance on physical habitat structures for predation.

### Defensive Mechanisms of Blackworm Blobs

Shifting from the dynamics between a single leech and an individual blackworm to a population of highly entangled worm blobs reveals a change in predation outcomes. In these complex assemblies, we observe defensive maneuvers not just by the targeted worm but by the collective, significantly impeding the leech’s predatory success.

When the leech places its anterior sucker on a worm within the blob, it encounters undulating movements as the worm, while remaining entwined with its conspecifics, vigorously resists (Fig.3a, t=0 to 0.5s). These movements draw the entire blob towards the predator (t=0.5 to 1s), rather than isolating the prey. Subsequent attempts by the leech to envelop the entire blob through its ventral fold are thwarted by the group’s collective density, presenting an insurmountable barrier and leading to a series of failed predation attempts (Fig.3a, t *>* 1.5s).

**Fig. 3.**
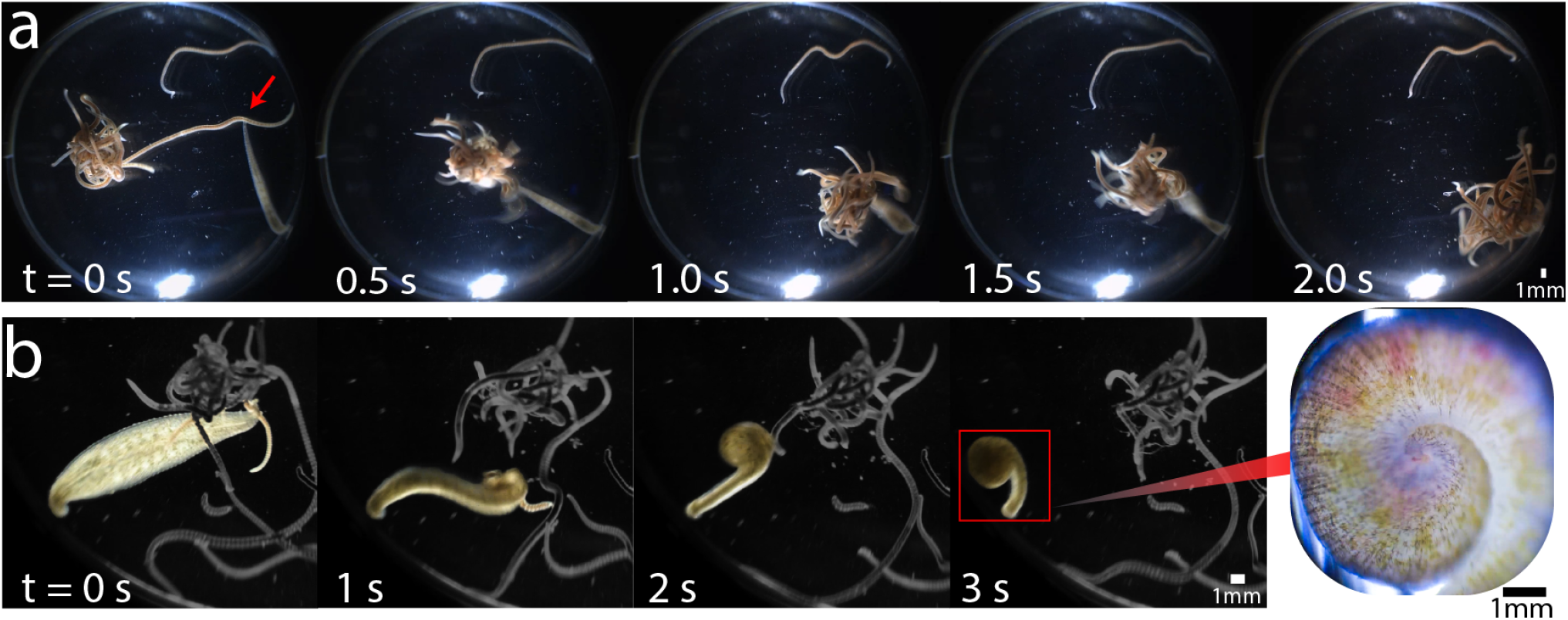
Predation by *Helobdella stagnalis* on worm blobs. **(a)** Timelapse of a leech’s failed attempt to trap an individual worm from a blob. After attaching its anterior sucker to a worm (red arrow), the worm undulates violently while maintaining itself entangled with conspecifics (t=0 to 0.5s). This causes the leech to pull the entire blob towards itself (t=0.5 to 1.0s). At t *>* 1.5s, the leech attempts to execute a ventral fold onto the blob several times but inevitably fails. The inset shows a worm completely trapped inside of the cavity. **(b)** Successful complete entombment of a worm from an entangled population of (*∼* 10 worms). (**Supplementary movie 3 and 4)**.

Fig.3b, captures a successful entombment by the leech, with a worm fully trapped within the formed cavity. Finally, as shown in Fig.3c, the leech inserts its proboscis to ingest the internal contents of the prey. These observations indicate that the worm blob’s defensive strategy, through collective undulation and cohesion, challenges the leech’s ability to isolate and entrap prey. This defense mechanism suggests an evolved collective response, perhaps an adaptation driven by persistent predatory pressures from leeches like *Helobdella* spp.

## Discussions

Our observations show that *Helobdella*, typically effective in feeding on less reactive prey like mollusks or *T. Tubifex*, employ a specialized entombment strategy when preying on blackworms. This adaptation allows the leeches to counter the blackworms’ escape capabilities and collective entanglement strategy. Unlike *T. Tubifex* worms from the Naididae family, which lack significant defensive reactions, blackworms exhibit robust escape behaviors, indicating their closer evolutionary proximity to leeches (Kuo and Lai 2019).

Abiotic factors significantly impact these predator-prey dynamics. The dense aggregation of blackworms in blobs may influence local oxygen levels, potentially affecting the respiratory efficiency of the leeches (Tuazon et al. 2022; Savoie et al. 2023; Deblais et al. 2023. *Helobdella* spp., which respire through their skin, sometimes supplement their oxygen intake through undulating movements, a behavior also seen post-feeding or in their parental care practices. During these activities, adult *Helobdella* spp. actively ventilate their young, enhancing the oxygen content around them, which is crucial in low-oxygen environments (Kutschera and Wirtz 2001). We hypothesize that this adaptation could be disadvantageous when a leech attempts to feed within a worm blob, as the collective might reduce the available oxygen, potentially disrupting a feeding leech’s ability to aerate (see Supplementary movie 5).

The habitat preferences of blackworms significantly contribute to their survival strategies. In addition to residing under detritus, studies show that blackworms often inhabit areas beneath macrophytes, such as duckweed (Fig. 1a). These plants provide physical shelter and enhanced nutrient conditions, potentially as a defensive adaptation against predators (Ohtaka et al. 2011, 2014; Xie et al. 2008; Vanamala Naidu et al. 1981; Cheruvelil et al. 2002). This “interrhizon” or the rhizosphere around plant roots facilitates better oxygenation, supporting dense worm populations that are less likely to be preyed upon by predators, such as *Helobdella*, which are less frequently found in vegetated areas (Ohtaka et al. 2011; Jab-loń ska-Barna 2007; Waters and San Giovanni 2002; Talbot and Ward 1987). This ecological setup highlights the critical role of habitat in shaping the interactions and survival strategies of both blackworms and their leech predators.

### Limitations and Future Outlook

Our study highlights the predator-prey dynamics between *Helobdella* and *Lumbriculus variegatus*, but it has some limitations. We relied on observational and experimental methods without extensive quantitative analysis, limiting our ability to statistically validate the ecological impacts. The broad identification of our experimental organisms as *Helobdella* spp. also poses a limitation. Without detailed molecular analysis, species-specific variations in prey selection and predatory strategies may influence the generalizability of our findings. Each species of liquidosomatophagous freshwater leech may exhibit unique behaviors, and the spiral entombment strategy observed may not apply universally across all *Helobdella* species.

Despite these limitations, our primary goal was to document and analyze the spiral entombment strategy. Our findings offer focused insights into this behavior in the context of *Helobdella* interactions with blackworms. Further research is needed to determine the prevalence and variation of this strategy among other leech species in different ecological settings.

## Conclusions

Our study described a unique entombment strategy used by *Helobdella* spp. when preying on blackworms. Our observations highlight a potentially adaptive response to the rapid escape and defensive mechanisms of blackworms, showcasing a complex interaction previously underexplored in aquatic ecosystems. This research could inspire future studies on the prevalence of this behavior across different leech species and its ecological implications. Future studies could explore how variations in environmental conditions, such as oxygenation and vegetation, and prey characteristics influence the evolution and efficacy of these predatory strategies.

## Supplementary data

Supplementary data available at *ICB* online.

## Author contributions statement

H.T. and M.S.B. conceptualized the research. H.T. and W.D. designed the experiments. H.T., and W.D conducted the experiments, for which H.T., W.D., K.M. performed the analysis. M.S.B. supervised the research. We thank Dr. Ishant Tiwari for assisting with the false coloring figure and Ivy Li for assisting with the rearing of the worms and duckweed. All authors contributed to writing, discussion, and revising the manuscript.

## Acknowledgments

H.T. acknowledges funding support from the NSF graduate research fellowship program (GRFP) and Georgia Tech’s President’s Fellowship. M.S.B. acknowledges funding support from NIH Grant R35GM142588; NSF Grants MCB-1817334; CMMI-2218382; CAREER IOS-1941933; PHY-2310691 and the Open Philanthropy Project. Text in this paper was revised using ChatGPT-4.

